# Safety, pharmacokinetics, and potential neurological interactions of ivermectin, tafenoquine and chloroquine in Rhesus Macaques

**DOI:** 10.1101/2024.02.01.578449

**Authors:** Pattaraporn Vanachayangkul, Chanikarn Kodchakorn, Winita Ta-aksorn, Rawiwan Im-erbsin, Anchalee Tungtaeng, Phornpimon Tipthara, Joel Tarning, Luis A. Lugo-Roman, Mariusz Wojnarski, Brian A. Vesely, Kevin C. Kobylinski

## Abstract

Ivermectin could be used for malaria control as treated persons are lethal to blood feeding *Anopheles*, resulting in reduced transmission. Tafenoquine could be used in combination with ivermectin to clear persons of liver stage *Plasmodium vivax* reservoir and as a prophylactic in high-risk populations. The safety of ivermectin and tafenoquine has not been evaluated. As earlier forms of 8-aminoquinolones were neurotoxic, and ivermectin is an inhibitor of the P-glycoprotein blood brain barrier transporter, there is concern that co-administration could be neurotoxic. The safety and pharmacokinetic interaction of tafenoquine, ivermectin, and chloroquine was evaluated in Rhesus macaques. No clinical, biochemistry, or hematological outcomes of concern were observed. The Cambridge Neuropsychological Test Automated Battery was employed to assess potential neurological deficits following drug administration. Some impairment was observed with tafenoquine alone and in the same monkeys with subsequent co-administrations. Co-administration of chloroquine and tafenoquine resulted in increased plasma exposure to tafenoquine. Urine concentrations of the 5,6 orthoquinone TQ metabolite were increased with co-administration of tafenoquine with ivermectin. There was an increase in ivermectin plasma exposure when co-administered with chloroquine. No interaction of tafenoquine on ivermectin was observed *in vitro*. Chloroquine and trace levels of ivermectin, but not tafenoquine, were observed in the cerebrospinal fluid. The 3”-*O*-demethyl ivermectin metabolite was observed in macaque plasma but not in urine or cerebrospinal fluid. Overall, the combination of ivermectin, tafenoquine, and chloroquine did not have clinical, neurological, or pharmacological interactions of concern in macaques, therefore this combination could be considered for evaluation in human trials.

## Introduction

Tafenoquine (TQ) is a member of the 8-aminoquinoline (8-AQ) class of antimalarials which was recently approved for the radical cure of relapsing forms of *Plasmodium vivax* hypnozoites. Primaquine (PQ), also an 8-AQ, has been in clinical use for treatment of *P. vivax* hypnozoites since the 1950s. The advantage of TQ over PQ is that TQ is administered as a single dose to achieve vivax radical cure (1), while PQ requires seven to fourteen days of daily dosing (2). Neither PQ nor TQ have been associated with neurological complications. However, earlier 8-AQ derivatives such as plasmocid, pamaquine, and pentaquine were neurotoxic, causing damage specific to neuro-anatomical structures in both Rhesus macaques and humans which led to deficits in neurological function (3). P-glycoprotein (P-gp) is an efflux pump found along the blood brain barrier (BBB) that removes the drugs from the central nervous system (CNS), preventing toxic levels in the CNS. One possible explanation for the lack of TQ neurotoxicity is active efflux by P-gp, however, it is not known if TQ is a substrate for P-gp transport (4). It is unclear if co-administration of drugs that inhibit P-gp could result in increased rates of neurotoxic adverse events post TQ administration.

Ivermectin (IVM) mass drug administration (MDA) is under consideration as a novel vector control tool to reduce malaria transmission as treated hosts are lethal to blood feeding *Anopheles*. Ivermectin MDAs in West Africa have been shown to kill wild African *An. gambiae* (5,6), reduce the proportion of *P. falciparum* infectious *An. gambiae* (6,7), shift mosquito population age structure (6), and reduce malaria incidence in humans (8). Clinical trials have demonstrated that IVM can be co-administered with antimalarial drugs such as artemether-lumefantrine (9), dihydroartemisinin-piperaquine (DP) (10,11), and single low dose PQ (10). Recent field evidence from The Gambia indicates that IVM and DP MDAs can be safely administered at scale to reduce *Plasmodium falciparum* incidence and prevalence (12).

With both TQ and PQ there is risk for hemolytic toxicity in G6PD-deficient (G6PDd) persons. However, even with this known G6PDd risk, PQ MDAs for *P. vivax* radical cure have been conducted in populations with varying degrees of G6PD frequency in numerous countries including Afghanistan, Azerbijan, Tajikistan, North Korea (13), Taiwan, Papua New Guinea, Solomon Islands, Tanzania, Nicaragua, Malaysia, Indonesia, China, Kyrgyzstan (14), Vanuatu (15), and Cambodia (16). Recent advances in point of care diagnostics for G6PD deficiency (17) make screening for this issue during MDAs possible, reducing risk of G6PD-associated hemolysis. Thus, it is possible that TQ MDAs for *P. vivax* radical cure could be considered in the future. Targeted prophylaxis of populations at high-risk for malaria with TQ is also under consideration.

Ivermectin and TQ could be co-administered during MDAs for *P. vivax* radical cure and suppression of *Plasmodium* transmission in mosquito populations. Ivermectin is a known substrate and inhibitor of P-gp (18–20), thus, if TQ is a P-gp substrate then there is potential that IVM could alter TQ concentrations in the CNS when co-administered. Therefore, it is important to evaluate potential IVM and TQ drug-drug interactions (DDIs), particularly on neurological function. While a published expert opinion suggests that DDIs at the human BBB due to efflux transporter (*i.e.* P-gp) inhibition by marketed drugs are unlikely (21), it is still worth investigating for TQ as it was registered after this position statement was published and because of the historical neurotoxicity of 8-AQ drugs. Ivermectin is primarily metabolized by CYP3A4, with minor contribution of CYP3A5 and CYP2C8 (22). TQ has not been shown to affect the pharmacokinetics of midazolam (CYP3A4) or chloroquine (CYP3A4 and CYP2C8) in healthy human volunteers (4), so it is unlikely that co-administration with IVM would increase the exposure of TQ. Chloroquine (CQ) is frequently co-administered with TQ for blood and liver-stage *P. vivax* radical cure. Ivermectin and CQ have been shown to be safe to co-administer in macaques and CQ did not affect the exposure to IVM (23), but the combination of IVM, CQ, and TQ has not been investigated previously.

Regarding neurotoxicity of historical 8-AQs, both humans and Rhesus macaques display similar neurological signs following plasmocid, pamaquine, and pentaquine administration and include loss of pupillary reflex, nystagmus, disturbed eye movements, loss of equilibrium control, loss of motor coordination, postural hypotension, and syncope. Thus, the macaque serves as an ideal animal model to investigate potential neurological complications following 8-AQ exposure (3). Therefore, two veterinary trials were performed in Rhesus macaques to evaluate the potential pharmacokinetic and neurological DDIs of IVM, TQ, and CQ.

## Results

### Human liver microsome assay

The *in vitro* human liver microsome results showed no substantial difference in metabolic rate when IVM was incubated alone or together with TQ (Figure 1). The metabolism rate of IVM was evaluated based on the normalized peak area (%) of IVM over a total of 2.5 hours of incubation. The slope (95% CI) of the linear regression was based on the elimination of IVM and estimated to 0.30 (0.28-032) min^-1^ when IVM was incubated alone and to 0.25 (0.24-0.26) min^-1^. This suggests that TQ will have no substantial effect on the metabolism of IVM when co-administered (*i.e.* <20% relative difference).

**Figure 1.**
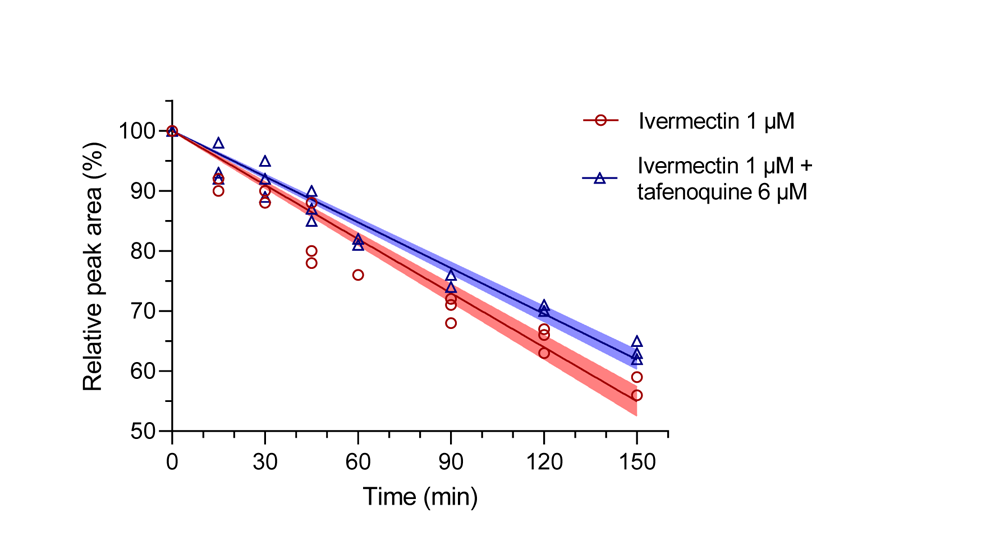
*In vitro* metabolism of ivermectin alone and when incubated with tafenoquine, using pooled human liver microsomes. Markers show peak-normalized IVM concentrations (three biological replicates per sampling time), the solid lines show a linear regression based on observed data, and the shaded areas show the 95% prediction interval of the linear regression analysis.

### Macaque Trial Safety Outcomes

#### Trial 1

No clinical neurological (*e.g.* ataxia, lethargy, imbalance) or gastroenterological (*e.g.* diarrhea, vomiting, weight loss) signs were observed in any of the macaques during Phases 1-3 drug administrations. All CBC parameters were within normal range. No blood urea nitrogen (BUN), total protein (TP), creatinine, or methemoglobin (MetHB) measurements were outside the range of normal. There were slight but not significant changes in macaque body weight. One of four macaques lost weight (2-3%) when provided TQ alone (Phase 1), two of four macaques lost weight (2-3%) and one macaque gained weight (5%) when provided TQ plus IVM (Phase 2), and finally three of four animals gained weight (1-3%) when provided TQ plus IVM plus CQ (Phase 3). One macaque had aspartate transaminase (AST) rising to grade 2 (R1441), and two macaques had AST rising to grade 1 (R1425, R1553) following TQ administration (Phase 1) and returning to baseline within 48 and 72 hours. One macaque had alanine aminotransferase (ALT) rising to grade 1 (R1441), following TQ administration (Phase 1) and did not return to baseline within 72 hours. No AST or ALT elevations above the upper limit of normal (ULN) occurred with TQ plus IVM (Phase 2) or TQ plus IVM plus CQ (Phase 3) (Figure 2). No total bilirubin or ALP measurements were above the ULN.

**Figure 2.**
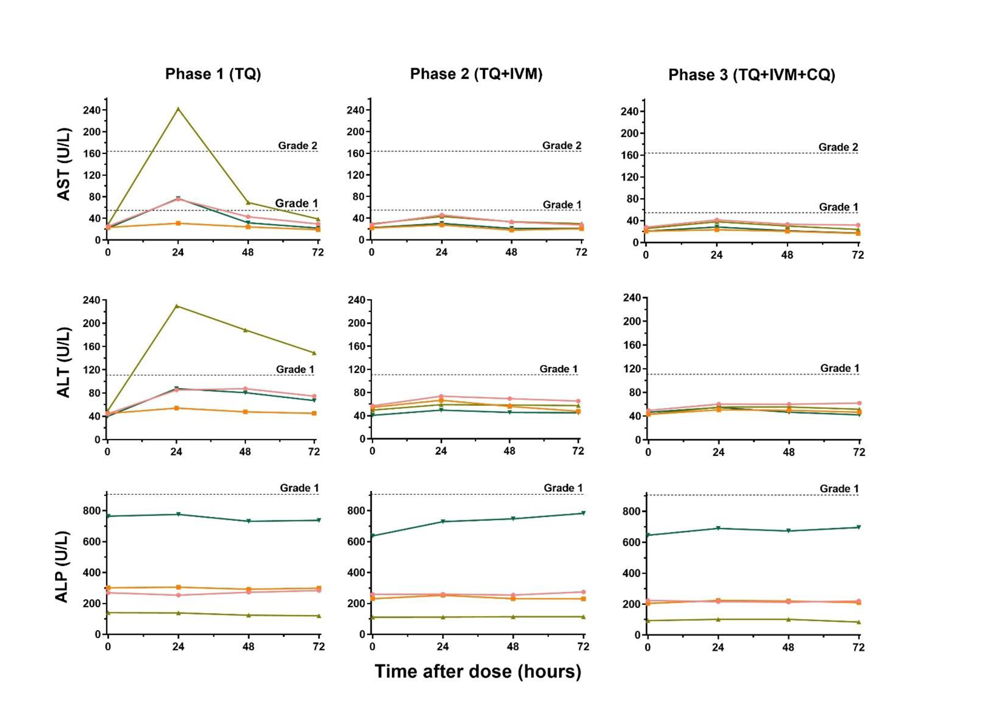
Liver function test results for all subjects, stratified by treatment regimen. Aspartate transaminase (AST), alanine aminotransferase (ALT), and alkaline phosphatase (ALP) values in all subjects following administration of tafenoquine (TQ) (Phase 1), TQ plus ivermectin (IVM) (Phase 2), or TQ plus IVM plus chloroquine (CQ) (Phase 3) at hours 0, 24, 48, 72 post treatment. Dashed lines represent grades above the upper limit of normal (ULN) for macaques housed at the Armed Forces Research Institute of Medical Sciences.

#### Trial 2

Again, no clinical neurological (*e.g.* ataxia, lethargy, imbalance) or gastroenterological (*e.g.* diarrhea, vomiting, weight loss) signs were observed in any of the macaques during Phases 4 and 5 drug administrations. As expected, transient mild to moderate reduced appetite was observed following anesthesia with ketamine for CSF collection, and appetite returned to normal after washout. There were slight but not significant changes in macaque body weight. One of four macaques lost weight (2%) when provided TQ plus CQ (Phase 4). Three of four macaques lost weight (1-2%) when provided TQ plus CQ plus IVM (Phase 5).

### Cambridge Neuropsychological Test Automated Battery (CANTAB) Results

#### Trial 1

No outlier results were observed for the Self-Ordered Spatial Search (SOSS) Accuracy Test. Outlier results indicating impairment were observed for the SOSS Duration Test for one macaque for TQ alone and the same macaque for TQ plus IVM combination (R1553). Outlier results indicating impairment were observed for the Motor Screening Task (MOT) Accuracy Test for three macaques (R1553, R1425, R1441) for TQ alone (Phase 1), and two of the same macaques (R1425, R1441) with TQ plus IVM (Phase 2), and again the same two macaques for TQ plus IVM plus CQ combination (Phase 3). Outlier results indicating impairment were observed for the MOT Number of Tests for three macaques (R1553, R1425, R1441) for TQ alone (Phase 1), the same three macaques with TQ plus IVM (Phase 2), and again for two of the same macaques (R1553, R1441) with TQ plus IVM plus CQ combination (Phase 3).

## Pharmacokinetics

### Trial 1

#### Plasma drug concentration-time profiles

Plasma TQ concentrations were increased following co-administration with CQ plus IVM (Phase 3), resulting in significantly higher exposure (AUC_∞_) compared to that seen for TQ administration alone (Phase 1) (p = 0.0126) and in combination with IVM (Phase 2) (p = 0.0219). Administration of IVM together with TQ did not cause any changes in TQ plasma profiles (Table 1; Figure 4).

**Table 1.** Pharmacokinetic parameters of tafenoquine in plasma when administered alone (Phase 1), with ivermectin (Phase 2), and with ivermectin plus chloroquine (Phase 3)

| PK Parameter | Tafenoquine |  |  |  |  |  |
| --- | --- | --- | --- | --- | --- | --- |
|  | TQ<br>(Phase 1)<br>Mean±SD | TQ+IVM<br>(Phase 2)<br>Mean±SD | TQ+IVM+CQ<br>(Phase 3)<br>Mean ± SD | P-value |  |  |
|  |  |  |  | Phase 1 v<br>Phase 2 | Phase 1 v<br>Phase 3 | Phase 2 v<br>Phase 3 |
| AUC<sub>%Extrap</sub> (%) | 2.3 ± 0.49 | 12.5 ± 3.31 | 11.9 ± 6.70 | **0.0065** | 0.7125 | 0.0618 |
| AUC<sub>∞</sub> (hr×ng/ml) | 3597 ± 1374 | 3248 ± 1583 | 6508 ± 1601 | 0.1889 | **0.0126** | **0.0219** |
| C<sub>max</sub> (ng/ml) | 67.4 ± 8.5 | 56.0 ± 18.6 | 84.4 ± 24.5 | 0.1688 | 0.2920 | 0.0865 |
| t<sub>1/2</sub> (hr) | 45.8 ± 10.3 | 49.4 ± 11.7 | 63.2 ± 11.7 | 0.4163 | **0.0130** | 0.1390 |
| T<sub>max</sub> (hr) | 12 ± 0 | 12 ± 0 | 12 ± 0 | N/A | N/A | N/A |
| CL/F (L/hr/kg) | 0.64 ± 0.31 | 0.75 ± 0.39 | 0.33 ± 0.10 | 0.1133 | 0.0618 | 0.0714 |
| Vz/F (L/kg) | 40.2 ± 13.9 | 49.0 ± 16.4 | 29.7 ± 9.72 | 0.1001 | **0.0464** | 0.0518 |

AUC<sub>%Extrap</sub> is the extrapolated area from AUC<sub>last</sub> to AUC<sub>∞</sub>, AUC<sub>∞</sub> is the total exposure, CL/F is the apparent elimination clearance, Vz/F is the apparent volume of distribution, C<sub>max</sub> is the maximum concentration, t<sub>1/2</sub> is the terminal elimination half-life, and T<sub>max</sub> is the time to reach the maximum concentration. Significant differences are in bold and red using one way ANOVA with Dunnet's method for multiple comparisons test.

Plasma IVM concentrations were higher when co-administering IVM plus CQ plus TQ (Phase 3) compared to IVM plus TQ (Phase 2), resulting in a significantly increased AUC_∞_ (p = 0.0165) (Table 2; Figure 5). Only the 3”-*O*-demethyl IVM metabolite was found in macaques at 4, 8,12 hours after dosing, when evaluating blood samples from TQ plus IVM (Phase 2) and TQ plus IVM plus CQ (Phase 3). The other two main metabolites found in humans, 4-hydroxymethyl IVM and 3”-*O*-demethyl, 4-hydroxymethyl IVM, were not measurable in the samples collected here. No novel ivermectin metabolite peaks were observed in the macaque plasma that were not previously observed in human plasma following a single oral dose at 400 µg/kg (22).

Plasma CQ and desethylchloroquine (dCQ) from IVM plus CQ plus TQ (Phase 3) reached C_max_ of 314.1 and 89.8 ng.ml at 7 and 11 hr respectively (Figure 6) (Table 3).

**Table 2.** Pharmacokinetic parameters of ivermectin in plasma when co-administered with tafenoquine (Phase 2) and tafenoquine plus chloroquine (Phase 3)

| PK Parameter | Ivermectin |  |  |
| --- | --- | --- | --- |
|  | TQ+IVM<br>(Phase 2)<br>Mean±SD | TQ+IVM+CQ<br>(Phase 3)<br>Mean ± SD | P-value |
| AUC<sub>%Extrap</sub> (%) | 10.5 ± 2.85 | 11.5 ± 1.59 | 0.6018 |
| AUC<sub>∞</sub> (hr×ng/ml) | 4349 ± 843 | 5493 ± 595 | **0.0165** |
| C<sub>max</sub> (ng/ml) | 153 ± 41.7 | 142 ± 9.90 | 0.5748 |
| t<sub>1/2</sub> (hr) | 22.4 ± 8.29 | 26.6 ± 6.57 | 0.3414 |
| $T_{\max}$ (hr) | $5.5 \pm 3$ | $7 \pm 2$ | 0.391 |
| CL/F (L/hr/kg) | $0.07 \pm 0.01$ | $0.06 \pm 0.01$ | **0.0405** |
| Vz/F (L/kg) | $2.18 \pm 0.46$ | $2.09 \pm 0.42$ | 0.7961 |

AUC<sub>%Extrap</sub> is the extrapolated area from AUC<sub>last</sub> to AUC<sub>∞</sub>, AUC<sub>∞</sub> is the total exposure, CL/F is the apparent elimination clearance, Vz/F is the apparent volume of distribution, C<sub>max</sub> is the maximum concentration, t<sub>1/2</sub> is the terminal elimination half-life, and T<sub>max</sub> is the time to reach the maximum concentration. Significant differences are in bold and red using Paired t-test.

**Table 3.** Pharmacokinetic parameters of chloroquine and desethylchloroquine in plasma when co-administered with tafenoquine plus ivermectin (Phase 3)

| PK Parameter | Chloroquine | Desethylchloroquine |
| --- | --- | --- |
|  | TQ+IVM+CQ<br>(Phase 3)<br>Mean ± SD | TQ+IVM+CQ<br>(Phase 3)<br>Mean ± SD |
| AUC<sub>%Extrap</sub> (%) | $4.82 \pm 2.25$ | $24.9 \pm 12.3$ |
| AUC<sub>∞</sub> (hr×ng/ml) | $7776 \pm 1801$ | $3553 \pm 991$ |
| C<sub>max</sub> (ng/ml) | $314 \pm 83.5$ | $89.8 \pm 30.4$ |
| t<sub>1/2</sub> (hr) | $19.5 \pm 0.50$ | $33.2 \pm 8.11$ |
| T<sub>max</sub> (hr) | $7 \pm 2$ | $11 \pm 8.87$ |
| CL/F (L/hr/kg) | $1.35 \pm 0.40$ | $3.03 \pm 1.05$ |
| Vz/F (L/kg) | $38.3 \pm 11.9$ | $146 \pm 60.5$ |

AUC<sub>%Extrap</sub> is the extrapolated area from AUC<sub>last</sub> to AUC<sub>∞</sub>, AUC<sub>∞</sub> is the total exposure, CL/F is the apparent elimination clearance, Vz/F is the apparent volume of distribution, C<sub>max</sub> is the maximum concentration, t<sub>1/2</sub> is the terminal elimination half-life, and T<sub>max</sub> is the time to reach the maximum concentration.

#### Urine drug concentration profiles

Co-administration of IVM with TQ (Phase 2) and with TQ plus CQ (Phase 3) showed higher C_max_ and AUC_0-24_ of 5,6 orthoquinone tafenoquine in urine than TQ alone (Figure 7; Table 4).

**Table 4.**
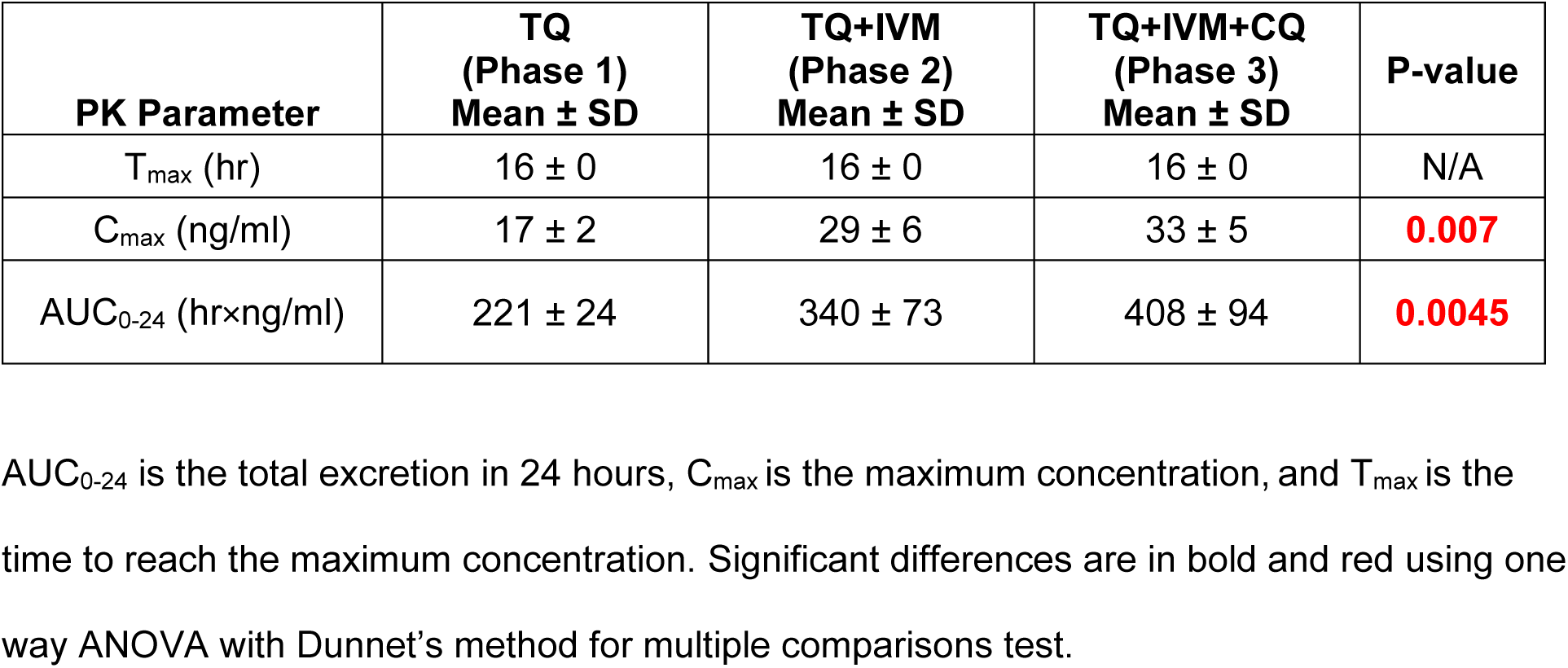
Pharmacokinetic parameters of 5,6 orthoquinone tafenoquine in urine with tafenoquine alone (Phase 1), co-administered with ivermectin (Phase 2), or ivermectin plus chloroquine (Phase 3)

| PK Parameter | TQ<br>(Phase 1)<br>Mean ± SD | TQ+IVM<br>(Phase 2)<br>Mean ± SD | TQ+IVM+CQ<br>(Phase 3)<br>Mean ± SD | P-value |
| --- | --- | --- | --- | --- |
| T<sub>max</sub> (hr) | 16 ± 0 | 16 ± 0 | 16 ± 0 | N/A |
| C<sub>max</sub> (ng/ml) | 17 ± 2 | 29 ± 6 | 33 ± 5 | **0.007** |
| AUC<sub>0-24</sub> (hr×ng/ml) | 221 ± 24 | 340 ± 73 | 408 ± 94 | **0.0045** |

AUC<sub>0-24</sub> is the total excretion in 24 hours, C<sub>max</sub> is the maximum concentration, and T<sub>max</sub> is the time to reach the maximum concentration. Significant differences are in bold and red using one way ANOVA with Dunnet's method for multiple comparisons test.

Ivermectin parent compound was detected in urine samples collected at 4, 8, 12 hours after dosing from TQ plus IVM (Phase 2) and TQ plus IVM plus CQ (Phase 3). Ivermectin was present in the 24 hour urine sample in the phase 3 samples, but at very low concentrations. No IVM metabolites were detected in urine.

### Trial 2

#### Blood and CSF Pharmacokinetics

Tafenoquine was detected in plasma but not CSF at 24 hours post-dose (Phases 4 and 5). The 5,6 orthoquinone TQ metabolite was not detected in plasma or CSF at 24 hours post-dose (Phases 4 and 5). Ivermectin was detected in plasma but not detected in CSF at 24 hours post-dose with the BEH method (Phase 5). Chloroquine and dCQ were detected in plasma and CSF at 24 hours post-dose (Phases 4 and 5) (Table 5).

**Table 5.**
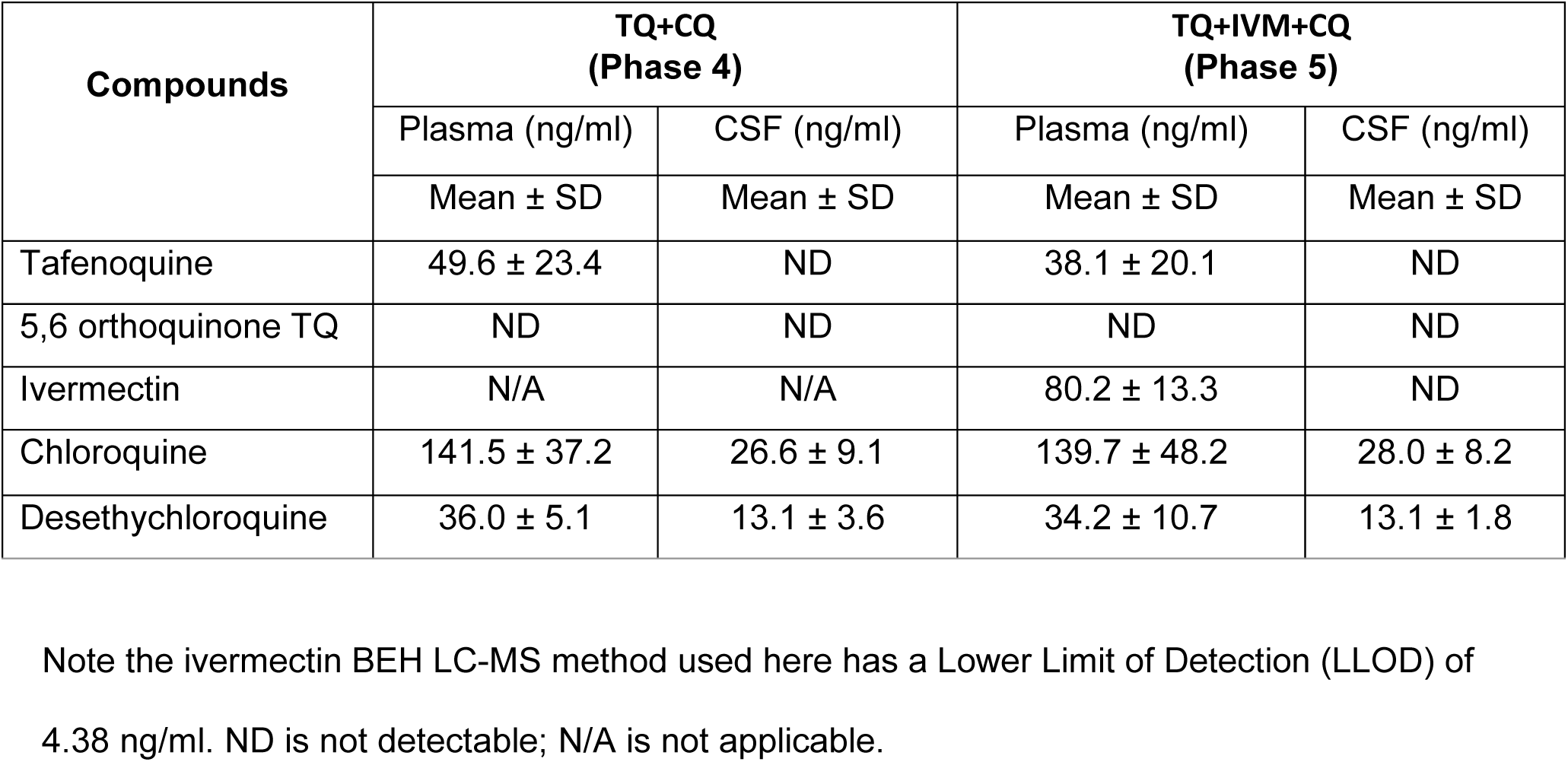
Tafenoquine, ivermectin, chloroquine, and desethylchloroquine concentrations in plasma and cerebral spinal fluid (CSF) 24 hours post-dose Note the ivermectin BEH LC-MS method used here has a Lower Limit of Detection (LLOD) of 4.38 ng/ml. ND is not detectable; N/A is not applicable.

#### Pharmacokinetic parameters of ivermectin and metabolites in CSF

Ivermectin parent compound was detected in CSF samples collected at 24 hours after dosing (Phase 5), but no IVM metabolites could be detected with the HSS T3 method.

## Discussion

There were no clinical neurological or gastroenterological symptoms noted in the macaques following any drug administrations from both trials (Phases 1-5). All CBC parameters, BUN, TP, creatinine, and MetHB results were within normal ranges during trial 1 (Phases 1-3). Three macaques had elevated AST following TQ alone (Phase 1), one rising to grade 2 and two rising to grade 1, all returned to baseline within 48 and 72 hours. One macaque had elevated ALT following TQ alone (Phase 1), rising to grade 1 and not returning to baseline within 72 hours. There were no increases in AST or ALT above the ULN for any drug combinations (Phases 2 or 3) (Figure 2).

There were some outlier CANTAB results following TQ alone (Phase 1) administration. However, any macaques with outlier CANTAB test results (SOSS, MOT) in drug combinations (Phases 2 and 3) were observed in the same macaques in TQ alone (Phase 1) administration (Figure 3). Thus, it does not appear that IVM or CQ coadministration with TQ led to further neurological impairment. The CANTAB impairment observed with TQ alone (Phase 1) is difficult to interpret as this was a small sample size (n = 4), no clinical signs of neurotoxicity were observed in this study, and previous macaque studies found no neurological concerns with TQ administration (3). To the best of our knowledge, this is the first study to use CANTAB to assess the acute effects of drug administration on macaque functionality. Further replication is necessary to determine if the SOSS and MOT impairment observed with TQ administration is repeatable in macaques.

**Figure 3.**
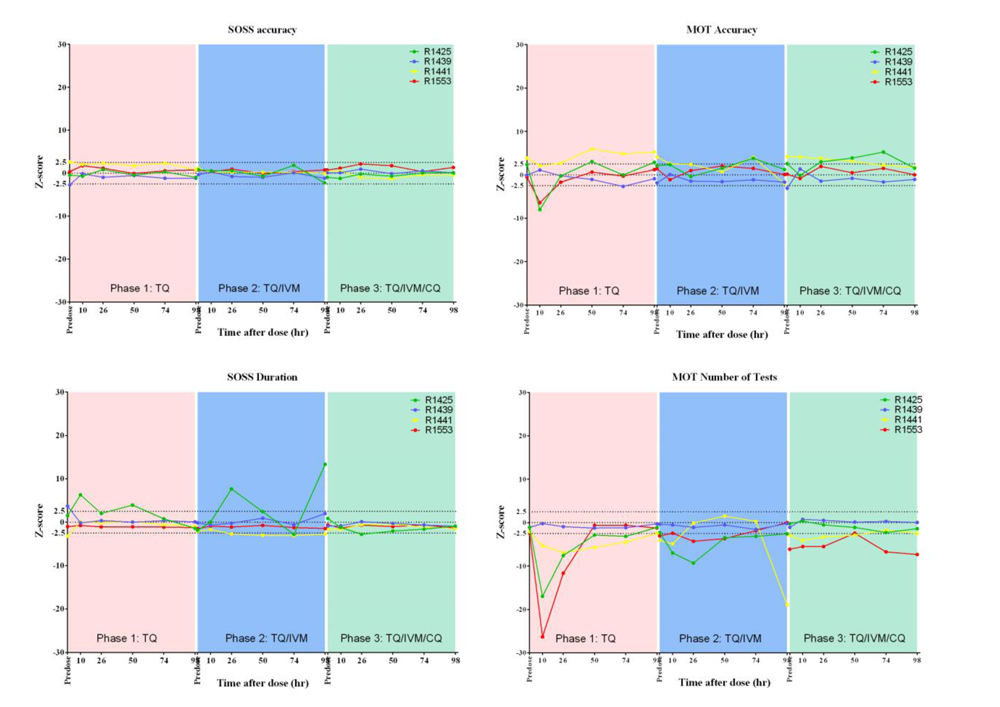
CANTAB results in macaques following administration of tafenoquine, ivermectin, and chloroquine combinations. Z-Scores for SOSS and MOT tests at hours 0, 10, 26, 50, 74, 98 for: (1) TQ alone (Phase 1; peach panels), (2) TQ plus IVM (Phase 2; blue panels), (3) TQ plus IVM plus CQ (Phase 3; green panels). The horizontal dashed lines indicate Z-scores 2.5 standard deviations from baseline normal, thus any values outside this range are considered outliers.

No TQ or 5,6 orthoquinone TQ metabolite was found in the CSF 24 hours post administration when co-administered with CQ (Phase 4) or with IVM plus CQ (Phase 5) (Table 5). Thus, it does not appear that IVM, at the dose administered, inhibits P-gp enough to allow TQ to penetrate beyond the BBB into the CNS. It is the opinion of the International Transporter Consortium that drug-drug induced neurotoxicity due to P-gp inhibition will be unlikely as there has to be enough unbound drug in circulation to achieve inhibition of P-gp greater than 90% (21). IVM has a 93% plasma protein binding (24) limiting the risk that unbound ivermectin in circulation is sufficient to saturate BBB P-gp, but the concentration of ivermectin needed to achieve saturation of BBB P-gp has never been established. Owing to the excellent safety profile of ivermectin after billions of treatments and in conjunction with the lack of reported CNS toxicities, supports the notion that BBB P-gp inhibition in humans is unlikely (21). These points suggest that IVM and TQ co-administration in humans would not lead to TQ penetrating beyond the BBB to a great extent. CQ was found in the macaque CSF (Phases 4 and 5) (Table 5), but CQ has been observed in human CSF with single dosing in malaria patients (25), thus presence of CQ in the CSF is not likely a result of a DDI caused by TQ or IVM.

Knockout mice lacking P-gp (26), CF-1 mice with P-gp deficiency (27), and collie dogs lacking P-gp due to a 4 bp deletion in the MDR1 gene (28) can experience neurological complications from IVM treatment at standard doses. In humans, there is one report of neurological consequence in a 13-year-old boy with scabies treated with oral IVM (230 µg/kg) that experienced coma, ataxia, pyramidal signs, and binocular diplopia. Interestingly, it was determined that there were mutations in his MDR1 gene which likely led to P-gp deficiency and subsequent adverse events following IVM treatment (29). Oral IVM has been used to treat nematode infections with CNS complications including *Baylisascaris procyonis* (30) and *Strongyloides stercoralis* (31). Reports of IVM evaluation in the CSF are limited to baylisascaris treatment of a 13 month old child wherein IVM (175 µg/kg) administration led to undetectable levels in CSF and blood levels of 8.6 ng/ml at 12 hours post administration (30), and in a comatose adult subject with chronic lymphocytic leukemia and disseminated strongyloidiasis treated daily with nasogastric IVM (300 µg/kg) and at day ten IVM blood levels 49.3 ng/ml and 0.14 ng/ml in CSF, suggesting less than 1% CNS penetrance (31).

In the present trial, IVM parent compound was detected in CSF 24 hours after co-administration with TQ and CQ (Phase 5) when using the HSS T3 LC-MS method but not the BEH LC-MS method. The samples for ivermectin and metabolite detection with the HSS T3 method were concentrated before injection to the LC-MS/MS leading to higher sensitivity. Additionally, the HSS T3 column resin differs from BEH in that the C18 column is longer so the retention time results in greater separation of the components meaning that HSS T3 method could detect a lower concentration of IVM. However, the HSS T3 LC-MS method was not used to quantify absolute concentrations of IVM. Since the IVM LLOD for the BEH LC-MS method is 4.38 ng/ml, the amount of ivermectin in CSF must be lower than this. To our knowledge this is the only report to have investigated CSF levels of IVM in macaques. Since there were no CSF evaluations in macaques treated with IVM alone it cannot be concluded whether this was caused by a DDI with TQ or CQ. Interestingly, at the BBB, P-gp secretes substrates into the blood, while at the blood-CSF barrier P-gp secretes substrates into the CSF (32). Furthermore, CSF IVM concentrations may not be predictive for CNS penetration as CSF concentrations are not representative of unbound brain concentrations of poorly permeable substrates for blood-CSF P-gp (21).

Human liver microsome results indicated that TQ co-administration does not influence the metabolism of IVM to great extent (Figure 1). Results showed <20% relative difference in the metabolic rate of IVM when incubated alone or together with TQ, suggesting no metabolic enzyme-related DDI, and it is therefore unlikely that co-administration of IVM and TQ would produce altered pharmacokinetic properties of IVM in patients. A previous study in macaques found that IVM (300 µg/kg) alone produced a similar C_max_ and 24-hours post-dose concentration in plasma (23) as observed here, suggesting no DDI interaction of IVM with TQ in macaques. Ivermectin was not administered alone, therefore it is not possible to evaluate the impact of TQ on the pharmacokinetic properties of IVM in this trial. Ivermectin exposure in blood was elevated when co-administrating IVM plus TQ plus CQ (Phase 3) compared to IVM plus TQ (Table 2; Figure 5), suggesting a DDI between IVM and CQ in these animals. Both compounds are predominantly metabolized by CYP3A4, suggesting that this DDI might be due to competitive enzyme inhibition. Moreover, CQ has also been shown to be a weak P-gp inhibitor, resulting in an increased exposure to other P-gp substrates when co-administered (33). Data collected here cannot discriminate between these possible mechanisms. However, a previous trial in macaques showed no interaction with IVM and CQ administration (23). This potential DDI needs to be evaluated further.

Tafenoquine exposure in blood was elevated by co-administration of CQ (Phase 3) but not IVM (Phase 2) (Table 1; Figure 4). The increased TQ exposure could be a result from a reduced elimination clearance of TQ when co-administered with CQ, but it is difficult to firmly establish this mechanism with so few animals in each group. TQ is metabolized by CYP2D6 (34) and CQ inhibits CYP2D6 activity (33) which may lead to a DDI. CQ is a weak inhibitor of P-gp (33), thus, if TQ is a P-gp substrate, then the increased exposure seen when co-administered with CQ could possibly be due to intestinal P-gp inhibition. However, previous healthy volunteer trial in humans showed no pharmacokinetic interaction between TQ and CQ (35). Thus, the DDI observed here requires further investigation. Conversely, IVM co-administration with TQ increased 5,6 orthoquinone TQ concentrations in urine (Phases 2 and 3) but CQ co-administration did not (Phase 3), but data collected here cannot be used to elaborate on a possible mechanism for this DDI (Table 4; Figure 7).

**Figure 4.**
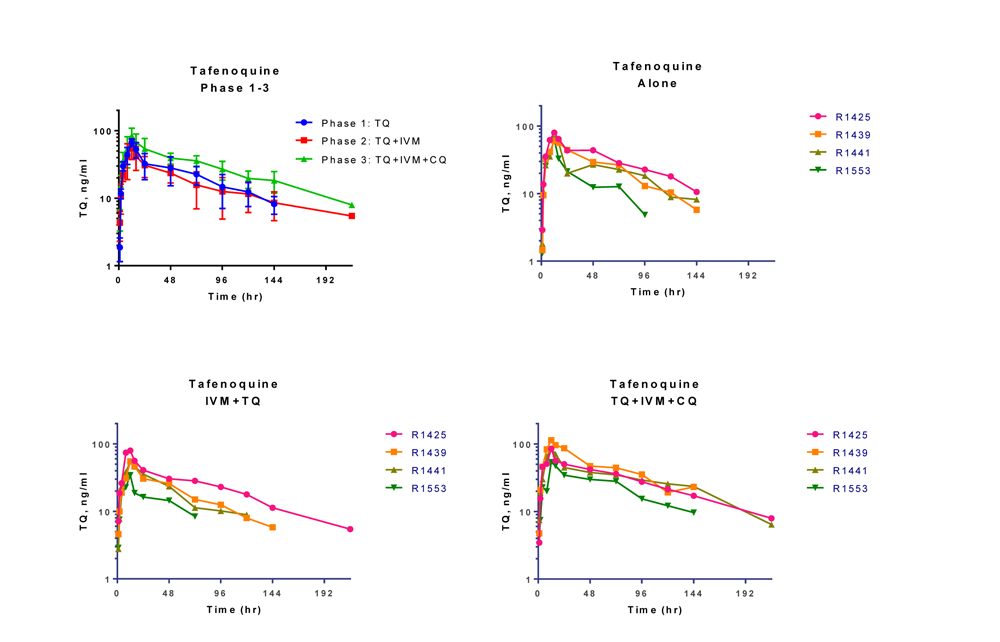
Plasma tafenoquine (TQ) concentration-time profiles in macaques. Tafenoquine plasma concentrations for (1) all subjects by Phases 1-3, (2) by subject for TQ alone (Phase 1), (3) by subject for TQ plus IVM (Phase 2), and (4) by subject for TQ plus IVM plus CQ (Phase 3).

**Figure 5.**
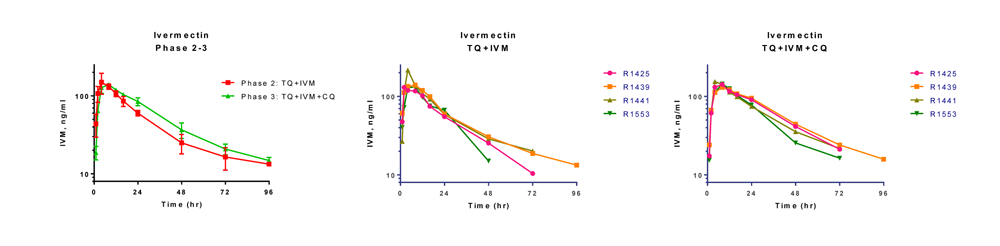
Plasma ivermectin concentration-time profiles in macaques. Ivermectin (IVM) plasma concentrations for (1) all subjects by Phases 2 and 3, (2) TQ plus IVM by subject (Phase 2), and (3) TQ plus IVM plus CQ by subject (Phase 3).

**Figure 6.**
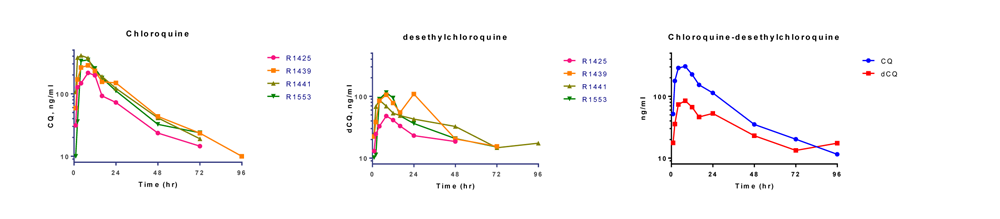
Plasma chloroquine (CQ) and desethylchloroquine (dCQ) concentration-time profiles in macaques. CQ plasma concentrations (1) by subject for TQ plus IVM plus CQ (Phase 3), (2) dCQ by subject for TQ plus IVM plus CQ (Phase 3), and (3) for all subjects CQ and dCQ from TQ plus IVM plus CQ (Phase 3).

**Figure 7.**
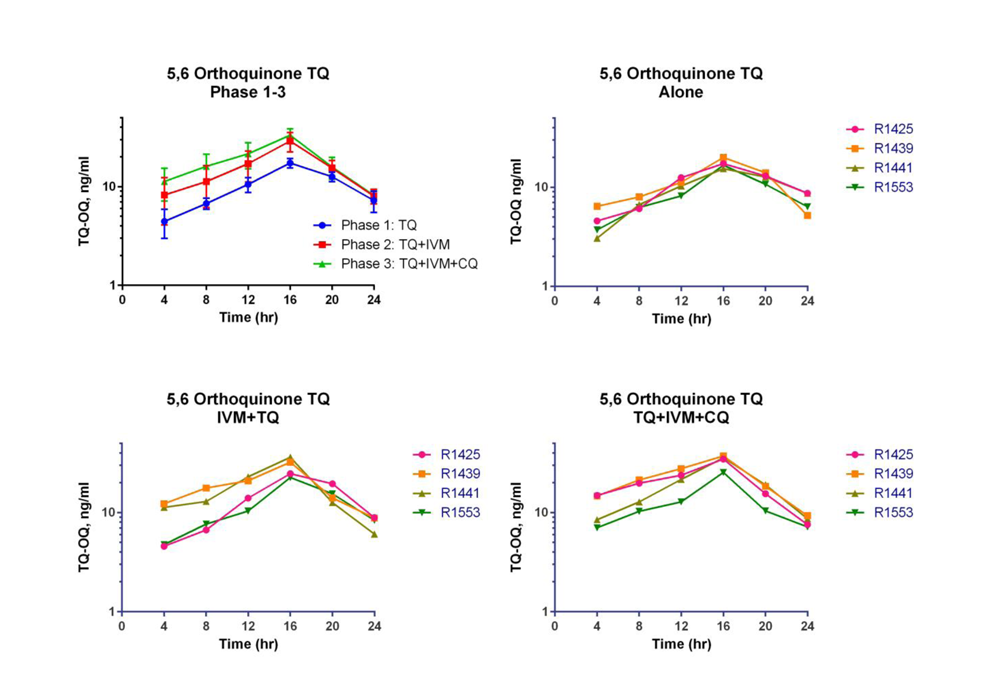
Urine 5,6 orthoquinone tafenoquine (TQ) concentration-time profile. 5,6 orthoquinone TQ urine concentrations for (1) all subjects by Phases 1-3, (2) TQ alone by subject (Phase 1), (3) TQ plus IVM by subject (Phase 2), and (4) TQ plus IVM plus CQ by subject (Phase 3).

Only one of the three primary IVM metabolites found in humans (22) was identified in macaque plasma, 3”-*O*-demethyl IVM metabolite. This indicates that IVM does not undergo hydroxylation at the C-4 carbon to produce the 4-hydroxymethyl IVM or 3”-*O*-demethyl, 4-hydroxymethyl IVM metabolites in macaques. No IVM metabolites were found in the CSF (Phase 5) or in urine (Phases 2 and 3). Ivermectin parent was identified in macaque urine with the HSS T3 method but the amount was not quantified. In humans the major route of IVM and metabolite elimination is fecal (36). Administration of IVM in cattle, sheep, swine, and rats (300 or 400 µg/kg) via subcutaneous, oral or intramural routes resulted in 0.5 – 2.0% of administered IVM in urine (37).

In conclusion, the safety, neurological, and pharmacological results presented here suggest that IVM and TQ combinations for MDAs or prophylaxis could be a viable option. However, the limited number of macaque subjects in this trial (n = 4), warrants further evaluation of potential DDIs which should be studied in a controlled human clinical trial.

## Materials & Methods

### Human liver microsome assay

#### Chemicals and materials

Ivermectin, IVM-D2, TQ, pooled human liver microsomes, β-Nicotinamide adenine dinucleotide 2′-phosphate reduced tetrasodium salt hydrate (β-NADPH), and ammonium acetate (LC-MS grade) were purchased from Sigma-Aldrich (St. Louis, MO, USA). Potassium phosphate 0.5 M buffer solution pH 7.4 was purchased from Thermo Fisher Scientific (Waltham, MA, USA). Ivermectin-D2 was purchased from Toronto Research Chemicals (Toronto, ON, Canada). Formic acid (LC-MS grade) was purchased from Honeywell Fluka (Seelze, Germany). LC-MS grade acetonitrile and methanol were purchased from J.T Baker (Phillipsburg, NJ, USA). Water (milli-Q 18.2 MΩ cm−1) was prepared from a Milli-Q purification system (Merck Darmstadt, Germany).

#### Preparation of drug solutions

Stock solutions of IVM (3.20 mM) were prepared in 80% acetonitrile, and TQ (3.95 mM) were prepared in 80% methanol and stored at −80°C until use. A working solution of IVM (100 µM) and TQ (600 µM) was prepared before use by dilution of the stock solution in the same solution. Ivermectin-D2 (100 ng/mL) was prepared in acetonitrile.

#### Microsomes assay

Pooled human liver microsomes (containing 20 mg/ml of protein) were thawed on ice. An aliquot of 0.1 M potassium phosphate buffer pH 7.4 (1,464 µL), 100 µM IVM (16 µL), and microsomes (40 µL) was mixed in a falcon tube kept on ice. The second tube was prepared by mixing aliquots of 0.1 M potassium phosphate buffer pH 7.4 (1,448µL), 100 µM IVM (16 µL), 600 µM TQ (16 µL), and microsomes (40 µL). A 470 µL aliquot of pre-mixed solution without TQ was transferred to a 96-well plate (Agilent Technologies, Santa Clara, USA) three wells (n = 3), and a 470 µL solution with TQ was transferred to another three wells (n = 3). The plate was incubated in a thermomixer at 37°C for 5 min with gentle shaking (300 rpm). After incubation, 20 mM NADPH (30 µL) was added and mixed well (total volume 500 µL). A 100 µL aliquot of sample was collected immediately to the new 96-well plate, each well containing 200 µL of ice-cold acetonitrile with 0.1 µg/mL of IVM-D2. The collecting plate was kept on ice. A 100 µL sample aliquot was collected at 15, 30, 45, 60, 90, 120, and 150 minutes. The collecting plate was sealed with the lid and centrifuged at 2100 g, 4°C for 45 min. A 100 µL aliquot of clear supernatant was transferred to the new 96-well plate, and LCMS analysis performed immediately.

#### Macaque Trials

Four healthy male macaques, 5 to 7 years old, and ranging in weight from 7.9 to 10.7 kg, with prior training for the Cambridge Neuropsychological Test Automated Battery (CANTAB) were selected for the study. All macaques were negative for simian retroviruses and simian herpes B virus. The macaques were included in two separate trials to evaluate potential DDIs of combinations of TQ, IVM, and CQ.

#### Trial 1

Trial 1 investigated plasma and urinary pharmacokinetics, and neurological evaluation of TQ alone (Phase 1), TQ plus IVM (Phase 2), and TQ plus IVM plus CQ (Phase 3). The Cambridge Neuropsychological Test Automated Battery (CANTAB) system was used to evaluate potential neurological impact of the study drugs. All study drugs were administered to pole-collar and chair restrained conscious macaques via nasogastric intubation at 1 ml/kg body weight. Monkeys were not anesthetized during nasogastric intubation. There was a 21-day washout period between each drug administration as the TQ half-life in macaques is 1-3 days (unpublished data).

For trial 1, just prior to drug administration and every 24 hours for 72 hours post-dose, venous blood was collected for evaluation of hematologic parameters were performed including methemoglobin (MetHB) and complete blood counts (CBCs), and liver parameters including alanine transferase (ALT), aspartate aminotransferase (AST), alkaline phosphatase (ALP), bilirubin, blood urea nitrogen (BUN), total protein (TP), creatinine. Blood for MetHB (0.5 ml) was collected in lithium heparin Vacutainer tubes. Blood for CBCs (0.5 ml) and biochemistry (0.5 ml) was collected in EDTA Vacutainer tubes. Any monkey that developed signs or symptoms of serious liver injury was withdrawn from the study. If elevated liver function results were observed then daily evaluation continued until return to baseline and results must be within normal range prior to initiating next drug treatment. Macaques were observed several times during the first few hours post dosing, and at least three times a day for the remainder of the study for any clinical signs of neurological (*e.g.* ataxia, lethargy, imbalance) or gastroenterological (*e.g.* diarrhea, vomiting, weight loss) complications.

#### Trial 2

The same four macaques were then evaluated to assess potential CNS penetration of study drugs following administration of: tafenoquine plus chloroquine (Phase 4), and TQ plus IVM plus CQ (Phase 5). CSF and blood were collected 24 hours before and after each drug administration and was assessed to determine the presence of the study drugs. For trial 2, no hematologic or liver parameters were assessed, no CANTAB evaluations were performed, and no urine collection for drug evaluation was performed. Macaques were observed several times during the first few hours post dosing, and at least three times a day for the remainder of the study for any clinical signs of neurological (*e.g.* ataxia, lethargy, imbalance) or gastroenterological (*e.g.* diarrhea, vomiting, weight loss) complications.

#### Drugs

Test compounds will be administered to unanesthetized macaques via nasogastric intubation. Only a single dose will be provided at each administration followed by a 21-day washout period. For trial 1, the drug regimens will be TQ alone (Phase 1), TQ plus IVM (Phase 2), and TQ plus IVM plus CQ (Phase 3). For trial 2, the drug regimens will be TQ plus CQ (Phase 4), and TQ plus IVM plus CQ (Phase 5). Study drugs will be administered at the following doses: TQ at 2 mg/kg, IVM at 0.3 mg/kg, and CQ at 10 mg/kg. Sparmectin-E (Sparhawk Laboratories, Inc., Lenexa, KS, USA) is a water-soluble formulation of IVM developed for oral use in horses that was diluted in sterile water. TQ succinate (WR238605-21) powder was dissolved in 25% Tween 80 solution and diluted in sterile water. CQ powder (Sigma Aldrich, Singapore) was dissolved and diluted in 0.5% hydroxyethylcellulose and 0.1% Tween 8 (HECT) solution.

#### Cambridge Neuropsychological Test Automated Battery (CANTAB) evaluation

For trial 1, the CANTAB system was used to assess neurocognitive deficits in macaques for testing memory, attention, and executive function by interacting with a touch-sensitive screen with food pellet reward. The Motor Screening Task (MOT) and the Self-Ordered Spatial Search (SOSS) Task were used to evaluate cognitive function following drug treatment. Because these tests measure latency (reaction speed) as a continuous variable, they are highly sensitive to detecting subtle changes in macaque neurocognition and therefore should be ideal for identifying potential neurocognitive deficits after treatment with the study drugs should they occur. The MOT assesses macaque response speed and accuracy by having a colored box appear in different locations on the screen and the macaque receives a food pellet reward when the box is selected, the MOT is run for ten minutes. The SOSS assesses macaque spatial memory by having several boxes appear on the screen with no obvious pattern and the animal is rewarded with a food pellet if it selects each box in turn without revisiting a box once it has been touched. Ten macaques were trained on the MOT and SOSS tests for 21 weeks prior to the initiation of the trial, and the four macaques with the highest and most consistent scores were selected for the trial. Prior to trial initiation, CANTAB MOT and SOSS evaluations were performed four times to establish a baseline score for each macaque. The MOT and SOSS evaluations were performed pre-dose and approximately 10, 26, 50, 74, 98 hours post-dose. For consistency, all CANTAB evaluations were performed approximately two hours after a venous blood collection occurred, including at baseline prior to drug administration.

Z-scores were used to evaluate the CANTAB results for each individual animal at each observation time point. The measurement of standard deviation below or above the mean for each animal as equation 1: Z = (x-µ)/σ. Variables defined as follows: Z: Z-score, x: baseline mean, µ: observed data, σ: baseline standard deviation. An impairment is defined when the Z-score is greater than 2.5 SD from the baseline mean. Statistical analyses were performed with GraphPad Prism version 8.0.

## Pharmacokinetics

### Venous sample collection

For trial 1, blood sampling (2 ml) for pharmacokinetics were performed at the following time points: at 0, 1, 2, 4, 8, 12, 16 hours and 1, 2, 3, 4, 5, 6, 10, and 14 days after each dose. For trial 2, blood sampling (2 ml) for pharmacokinetics was performed at 24 hours before and after each dose. Blood was collected in lithium heparin Vacutainer tubes and centrifuged at 3,000 rpm for 10 min and then the supernatant (plasma) was removed and kept at −80°C until analysis was performed. Plasma was separated into two tubes with 100-250 µl in each tube.

### Urine sample collection

For trial 1, urine was collected to determine orthoquinone presence/absence and approximate urine levels of the more polar tafenoquine orthoquinone which has been observed in human urine. Macaques were individually housed during the study and a tray was placed underneath each cage to collect urine. The cage trays were collected every 4 hours for 24 hours after dosing. The volume of urine excreted, urine creatinine and specific gravity was determined for each four-hourly urine collection sample. Approximately 5-10 ml of urine was collected from the tray at each collection, dispersed to cryopreservation tubes and kept at −80°C until analysis was performed.

### Cerebral Spinal Fluid (CSF) sample collection

For trial 2, cerebral spinal fluid (CSF) was collected 24 hours before and after each dose. Ketamine (15-20 mg/kg) was administered via intramuscular injection to anesthetize the macaques prior to CSF collection. CSF (1 ml) was collected via lumbar puncture with a 21 gauge, 3.8 cm needle. The CSF was dispersed to a cryopreservation tube and kept at −80°C until analysis was performed.

### Compound Extraction (BEH method)

#### Plasma and CSF sample preparation

100 µL of plasma and 50 µl of CSF were placed in 1.5 mL micro-centrifuge tubes, followed by protein precipitation by addition of 200 and 100 µL of acetonitrile with internal standard (SIL-PQ for TQ, IVM-D2 for IVM and MQ for chloroquine) for plasma and CSF, respectively. After 1.0 min of mixing, the samples were centrifuged at 9391*g* for 10 min, and then the 200 µL and 100 µl of supernatant from plasma and CSF, respectively, was filtrated through 0.22 µm PTFE membranes. The samples were then transferred to blue-cap vials and 5 µL of samples were injected into the UPLC-QToF system.

#### Urine sample preparation

200 µL of urine was placed in 1.5 mL micro-centrifuge tubes, followed by protein precipitation by addition of 400 µL of acetonitrile with internal standard (SIL-PQ for tafenoquine, IVM-D2 for IVM) and 400 µl of acetonitrile: methanol (80:20 v/v) with internal standard (MQ for chloroquine). After 1.0 min of mixing, the samples were centrifuged at 9391*g* for 10 min, and then the 400 µL of supernatant from urine, was filtrated through Clean Screen FASt® columns. The samples were then transferred to blue-cap vials and 5 µL of samples were injected into the UPLC-QToF system.

#### Liquid chromatography-mass spectrometry analysis (BEH method)

The liquid chromatography-mass spectrometry (LC-MS) was performed on Waters Acquity UPLC^TM^ equipped with Waters Xevo® G2-XS QToF (Waters Corp., Milford, MA, USA) operating in multiple reaction monitoring (MRM) mode. Chromatographic separation was performed on a Waters Acquity UPLC^®^ BEH C18 column (50x2.1 mm, 1.7 µm particle size) protected by a C18 guard column.

#### Tafenoquine (and metabolites)

The LC conditions consisted of a 5 minutes gradient of 5 mM ammonium acetate pH 4.5 in water (mobile phase A) and acetonitrile (mobile phase B). The LC column temperature was set to 40°C, and the auto-sampler temperature was set at 10°C. The injection volume was 5 µl. The linear gradient elution method was as follows: initial starting at 5% B (0.0-1.0 min), followed by increasing to 98% B (1.0-3.5 min), remaining at 98% B (3.5-4.0 min), decreasing to 5% B (4.0-4.5 min) and retained until the next sample injection (4.5-5.0 min). MS conditions were optimized in the positive electrospray mode with at capillary voltage 3.00 kV, source and desolvation temperature at 150 and 600°C, respectively, cone and desolvation gas flow at 50 and 600 L/h, respectively. The collision energies varied between 15 to 30 eV. The m/z of TQ, 5,6 orthoquinone TQ, and SIL-PQ (IS) were 464.3748 to 379.2530, 304.2710 to 191.1477 and 179.1120 to 247.1211, respectively. The MassLynx™ software (version 4.20, Waters Corp., Milford, MA, USA) were used for data acquisition and processing. The lower limits of quantification (LLOQ) were 3.99 and 10.0 ng/mL for TQ and 5,6 orthoquinone TQ, respectively in plasma, CSF, and urine.

#### Ivermectin

A gradient mode of mobile phase A (5mM ammonium acetate in water + 0.1% formic acid) and mobile phase B (5mM ammonium acetate in methanol + 0.1% formic acid) with column temperature of 40°C, at flow rate 0.40 mL/min with the injection volume was 5.0 µL as follows: initial starting at 50% B, and then increasing to 95% B within 2.0 min, remaining at this percentage for 4.0 min, after which, it was decreased to 50% B within 0.5 min and retained for 0.5 min. The total run time was 7 min and the injection volume was 5 µl. For mass spectrometry was set in the positive electrospray ionization mode with multiple reaction monitoring. Instrument parameters included capillary voltage of 3.5 kv, source and desolvation temperature of 150 and 400°C, respectively. The nitrogen generator was set at 120 lb/in^2^ to generate cone and desolvation gas flow of 100 and 800 L/H, respectively. The mass transitions were observed at m/z 892.77 è 569.50 and 894.79 è 571.52 for IVM and IVM-D2, respectively. Masslynx™ software (Waters Corp., Milford, MA, USA) was used for quantification. The IVM LLOQ for the BEH method was 10.0 ng/mL in plasma, CSF, and urine.

### Chloroquine

The gradient mobile phase of 5mM ammonium acetate pH 4.5 in water (mobile phase A) and acetonitrile: methanol (50:50, v/v) (mobile phase B) was used. The column temperature was at 40°C, auto-sampler temperature at 10°C and flow rate of 0.40 mL/min. The injection volume was 5 µL and total chromatographic run time was 5.0 minutes. The linear gradient elution method was as follows: initial starting at 5% B (0.0-1.5 min), followed by increasing to 75% B (1.5-3.0 min), decreasing to 5% B (3.0-4.0 min) and retained until the next sample injection (4.0-5.0 min). Mass Spectrometry was operated in positive ion and ion source ESI (electrospray ionization). The parameters were as follows: capillary voltage, 3.0 kV for positive mode: source temperature, 150 °C; desolvation temperature, 600 °C; cone gas flow, 100 L/h and desolvation gas flow, 1000 L/h. The collision energies varied between 10 to 42 eV. The multiple reaction monitoring (MRM) mode was used to monitor the transition of chloroquine m/z 320.33 è142.22, and mefloquine (internal standard) m/z 379.29 è 271.22, The MassLynx™ Software (Waters Crop., MS, USA) was used to control the instruments and chromatographic, peak integration and quantification. The MassLynx™ software (version 4.20, Waters Corp., Milford, MA, USA) were used for data acquisition and processing. The LLOQ was 9.30 and, 13.46 ng/mL for CQ and dCQ respectively in plasma and CSF.

### Compound Extraction (HSS T3 method) for ivermectin and metabolites

Plasma, CSF, and urine samples were thawed at ambient temperature. After vortexing and brief spin down, 50 µL of aliquots were transferred into a new microcentrifuge tube and mixed with 150 μL of cold acetonitrile. The tubes were put on Thermomixer and shaken for 10 min at 4°C. The sample tubes were centrifuged at 10,000*g* for 10 min at 4°C. The sample was concentrated by transferring 150 µL of supernatant into a new microcentrifuge and evaporated in a speed vacuum. The dried supernatant from plasma, CSF, and urine were reconstituted in 50 μL of 50% acetonitrile containing 0.1% formic acid and transferred to a LC vial for injection.

### Liquid chromatography-mass spectrometry (LC-MS) (HSS T3 method) for ivermectin and metabolites

The LC system was an ultra-high performance liquid chromatography (UHPLC) consisting of a quaternary LC pump (Agilent 1260), a high performance autosampler (Agilent 1260) set at 6°C, and a thermostatted column compartment (Agilent 1290) set at 40°C (Agilent Technologies, Santa Clara, USA). Five microliters of samples were analysed on a reversed-phase column ACQUITY UPLC HSS T3 (2.1 × 100 mm, 1.8 µm) protected by a precolumn ACQUITY UPLC HSS T3 (2.1 × 5 mm, 1.8 µm) (Waters Corporation, MA, USA) under linear gradient conditions using a mobile phase of water containing 10 mM ammonium acetate and 0.1% formic acid (mobile phase A) and acetonitrile:water (95:5, v/v) containing 10 mM ammonium acetate and 0.1% formic acid (mobile phase B) at a flowrate of 0.3 ml/min. The LC gradient started at 75% B and ramped to 90% B in 2 min (0-2 min). The gradient held constant at 90% B for 4 min (2-6 min). The gradient returned to 75% B in 0.1 min (6-6.1 min) with 3.9 min re-equilibration (6-10 min) until the next injection. A TripleTOF 5600+ quadrupole time-of-flight mass spectrometer; Q-TOF MS (Sciex, MA, USA), with a DuoSpray ionization source interface operated for electrospray ionization (ESI), was used for LC-MS analysis. The MS was operated in ESI-positive mode at ion source gas 1 (GS1) of 40 psi, ion source gas 2 (GS2) of 40 psi, curtain gas (CUR) of 30 psi, ion spray voltage floating (ISVF) of 4500 V, source temperature (TEM) at 350°C, and declustering potential (DP) of 120 V. Data were acquired in the TOF-MS scan at the mass range of *m*/*z* 100–1000. Data acquisition was performed using Analyst^®^ TF Software 1.8 and quantification of IVM level was performed by MultiQuant^TM^ Software (Sciex, MA, USA) with high resolution TOF-MS scan mode.

### Pharmacokinetic analysis

#### Plasma pharmacokinetics

Noncompartmental analysis (NCA) with linear trapezoidal-linear interpolation was used to generate pharmacokinetic parameters for IVM, TQ, and CQ using Phoenix WinNonlin 8.1 (Certara USA, Inc., NJ, USA). The pharmacokinetic parameters determined were the elimination half-life (T_1/2_), maximum concentration in plasma (C_max_), time to reach C_max_ after dosing (T_max_), area under the concentration-time curve until the last sample collection (AUC_last_), area under the concentration-time curve after the last dose to infinity (AUC_INF_) and percentage of AUC_INF_ due to extrapolation from T_last_ (last collection time point) to infinity (AUC_%Extrap_) and since the fraction of dose absorbed cannot be estimated for extravascular models, apparent volume of distribution (Vz/F) and apparent clearance (CL/F) were substituted for V and CL. Concentration data that lower than LLOQ were excluded. Statistical analysis and graphical representation were completed using GraphPad Prism version 6.0. One-way ANOVA with multiple comparison of Dunnett were used

#### Urine pharmacokinetics

Maximum concentration in urine (C_max_), time to reach C_max_ after dosing (T_max_) were observed and area under the concentration-time curve in 24 hour (AUC_24hr_) was calculated using GraphPad Prism Version 6.0

#### Ethical Statement

The USAMD-AFRIMS Institutional Animal Care and Use Committee and the Animal Use Review Division, U.S. Army Medical Research and Materiel Command, reviewed and approved this study (PN 20-05). Animals were maintained in accordance with established principles under the Guide for the Care and Use of Laboratory Animals eighth edition (38), the Animals for Scientific Purposes Act (39) and its subsequent regulations (40). The USAMD-AFRIMS animal care and use program is fully accredited by the Association for Assessment and Accreditation for Laboratory Animal Care (AAALAC). Following the Guide (38), animals enrolled in this study were part of the environmental enrichment program which aims to enhance animal well-being by providing the macaques with sensory and motor stimulation to facilitate the expression of species-typical behaviors and promote psychological well-being. All macaques were housed under conditions that provide sufficient space in accordance to established rules and regulations. Macaques were housed individually, however, opportunity for direct and indirect contact with conspecifics was provided to maintain their social environment. Animal care and husbandry were provided throughout the duration of the study by trained personnel and under the direction of licensed veterinarians.

## Acknowledgements

We thank the AFRIMS Department of Veterinary Medicine for conducting the macaque trial especially Laksanee Inamnuay, Kesara Chumpolkulwong, Natthasorn Komchareon, Chardchai Burom, Noppon Popruk, Sujitra Tayamun, Arasa Suttana, Mana Saithasao, Alongkorn Hanrujirakomjorn, Khrongsak Saengpha, Phakorn Wilaisri, Chakkapat Detpattanan, Rachata Jecksaeng, Siwakorn Sirisrisopa, Yongyuth Kongkaew, Sakda Wongsawanonkul, Sonchai Jansuwan, Amnart Andaeng, Chaisit Pornkhunviwat, Manas Kaewsurind, Wuthichai Puenchompu, Kidanan Rungrueang, Wuttichai Sawangsang, Dejmongkol Onchompoo, Paitoon Hintong, Siwadol Samano. Drs. Denise Hsu, Wanwisa Promsote, and Decha Silsorn (AFRIMS Dept of Retrovirology) for guidance on CANTAB data analysis. Funding was provided by the Military Infectious Disease Research Program. We thank Rattawan Kullasakboonsri for assistance with BEH method extractions and LC-MS/MS. The funders had no role in study design, data collection and interpretation, or the decision to submit the work for publication.

## Disclaimer

Material has been reviewed by the Walter Reed Army Institute of Research. There is no objection to its presentation and/or publication. The opinions or assertions contained herein are the private views of the author, and are not to be construed as official, or as reflecting true views of the Department of the Army or the Department of Defense. Research was conducted under an IACUC-approved animal use protocol in an AAALAC-accredited facility with a Public Health Services Animal Welfare Assurance and in compliance with the Animal Welfare Act and other federal statutes and regulations relating to laboratory animals

